# SpaGRN: investigating spatially informed regulatory paths for spatially resolved transcriptomics data

**DOI:** 10.1101/2023.11.19.567673

**Authors:** Yao Li, Xiaobin Liu, Lidong Guo, Kai Han, Shuangsang Fang, Xinjiang Wan, Xun Xu, Guangyi Fan, Mengyang Xu

**Affiliations:** BGI Research, Qingdao 266555, China; College of Life Sciences, University of Chinese Academy of Sciences, Beijing 100049, China; BGI Research, Beijing 102601, China; BGI Research, Shenzhen 518083, China; CAS Key Laboratory of Marine Ecology and Environmental Sciences, and Center of Deep Sea Research, Institute of Oceanology, Chinese Academy of Sciences, Qingdao 266071, China; BGI Research, Wuhan 430074, China; State Key Laboratory of Agricultural Genomics, BGI Research, Shenzhen 518083, China

**Author notes:** These authors contributed equally (Yao Li, Xiaobin Liu, Lidong Guo). Correspondence (G.F.), (M.X.).

## Abstract

Cells expressing similar transcriptional regulatory circuits spatially aggregate into distinct cell types or states. However, most existing methods for inferring gene regulatory networks from spatially resolved transcriptomics are devoted to spatial co-expression modules or interactions between transcription factors and target genes, neglecting mediated effects from extracellular signals. Here we introduce SpaGRN, a statistical framework for predicting the comprehensive intracellular regulatory network underlying spatial patterns by integrating spatial expression profiles with prior knowledge on regulatory relationships and signaling paths. We validate and assess SpaGRN using simulated and real datasets, demonstrating its efficiency, performance, and robustness. When applied to 3D datasets of developing *Drosophila* embryos and larvae, SpaGRN identifies spatiotemporal variations in specific regulatory patterns, delineating the cascade of events from receptor stimulation to downstream transcription factors and targets, revealing synergetic regulation mechanism during organogenesis. Moreover, SpaGRN provides flexible visualization functions. We construct an online 3D regulatory network atlas database for interactive exploration and sharing.

## INTRODUCTION

Studying the architecture of seemingly simple organisms or single tissues is an intricate undertaking that necessitates not only decoding the molecular profiles of numerous cells, but also comprehending the impact of their spatial context on cellular state and function^1^. Understanding the gene regulatory network (GRN) is imperative to decipher how cellular identity is established, maintained, and responds to the microenvironment at the molecular level^2^. The GRN represents a complex web of interactions between genetic materials including transcription factors (TFs) and downstream genes across different cell types and developmental stages. Although the GRN orchestrates cell functions via intracellular signaling within a cell, it is profoundly influenced by extracellular communications from neighboring cells in the spatial context^3,4^. For instance, the determination of distinct mesodermal cell fates in sea urchin embryonic development is driven by the spatially confined activation of the mesoderm GRN, which is restricted by the proximal endoderm cells^5^. Another compelling example is the remarkable insight gained from investigating the spatiotemporal behavior of histologically specific GRNs during *Drosophila* embryogenesis^6^.

Reconstructing GRNs is a fundamental goal of system biology leveraging diverse experimental methods and computational algorithms^7,8^. In this study, our focus is specifically on elucidating the interplay between TFs and target genes, with a particular emphasis on unraveling the intercellular effects through transcriptomics data. GRNs are mathematically represented as networks or graphs, which can be directed or undirected, weighted or unweighted, unipartite or bipartite, depending on the regulatory direction, strength, and gene sets under analysis. Pioneering efforts have leveraged bulk transcriptomics data to model GRNs by identifying gene co-expression patterns, which reflect the overall state of the tissue. For instance, WGCNA^9^ is a well-known correlation-based inference algorithm for predicting non-directional gene modules. However, this unsupervised approach is limited by the lack of regulatory relationships and cellular specificity, which can lead to false positive connections and hinder interpretability and causality.

Whereafter, GENIE3^10^ and its successor GRNBoost2^11^ employ random forest models trained for each target gene against all TF, incorporating prior regulatory information from TF–DNA binding assays, such as chromatin immunoprecipitation sequencing (ChIP-seq) experiments. The variable importance measures in these models uncover the regulatory potentials of TFs for targets, thereby introducing directional information to the inferred regulatory relationships. Meantime, the advent of single-cell technologies, particularly single-cell RNA sequencing (scRNA-seq), has opened up an entirely new avenue for GRN inference at the resolution of individual cells, enabling the tracing of cell cluster’s heterogeneity in GRNs across developmental stages or conditions^12–14^. scRNA-seq-based GRN inference algorithms primarily concentrate on exploiting cell type or state-specific regulatory patterns through linear and nonlinear regression analyses. This approach has facilitated temporal GRN modeling^15^, GRN perturbation analysis^16^, and GRN refinement by integrating other modalities such as scATAC-seq data^17^. Despite these advancements, predicting essential regulators to study cell-identity-specific phenomena using computational methods remains a challenging problem in bioinformatics^2,8^. For example, GRNs are essentially intrinsic to individual cells. But owing to the sparse and noisy nature, GRN inference algorithms for scRNA-seq data generally compute statistically significant co-expression of TF and targets for a specific cell cluster rather than one single cell. These algorithms assume that cells of the same type or state share the same regulatory events, neglecting the importance of considering the spatial context of TF-target co-expression. However, harnessing the co-expression information of neighboring cells can aid in the accurate inference of the GRN of a focal cell. Additionally, scRNA-seq-based GRN inference methods fail to account for the loss of cellular niche topological structure during cell dissociation, which can lead to the rewiring of the GRN inside a central cell due to ligand-receptor (L-R) interactions with spatially proximal cells. Thus, the connection between extracellular signaling and intracellular regulation is overlooked by these methods.

To address these challenges, we propose a refining and improving existing GRN model through spatially resolved transcriptomics (SRT). Recent advances in SRT technologies have enabled the investigation of spatially specific gene expression patterns and interactions within complex tissues^18–21^. Specifically, SRT allows for not only the inspection of gene co-expression pattern in specific spatial locations but also the identification of extracellular signaling that drives cellular responses from their respective niches. Current approaches to predict gene regulatory interactions using SRT data can be mainly categorized into four types: (a) subcellular proximity: extracting gene proximity relationships from subcellular patches to infer interaction networks using fluorescence *in situ* hybridization (FISH) imaging-based SRT data^22^; (b) direct application: directly using algorithms originally designed for scRNA-seq data to infer GRNs from SRT data^23^; (c) spatial co-expression: deriving spatially specific gene co-expression modules including graph-based Hotspot^24^, spatially weighted CellTrek toolkit^25^, and Bayesian-based SpaceX^26^; and (d) extracellular to intracellular model: exploiting the effect of cell-cell communications (CCC) to refine GRNs^27–29^. However, these methods pose new challenges owing to SRT data characteristics. In particular, the “subcellular proximity” strategy is only applicable to imaging-based ISH techniques to generate RNA profiles at the single molecule resolution for a limited number of preselected genes. The “direct application” approach allows for the GRN analysis of SRT data generated by *in situ* spatial barcoding-based platforms such as Stereo-seq, 10X Visium, and Slide-seq2. But challenges arise due to difficulties in precise cell segmentation, which may result from the presence of mixed cells within the same spot or the loss of pairwise ssDNA and H&E staining images. Moreover, the lack of spatial information in this approach can confound the identification of important TF-target associations^26^. Another dilemma is that most available methods cannot handle over tens of thousands of cells with tens of thousands of genes^8^. One compromise solution is cell aggregation or downsampling. Furthermore, the “spatial co-expression” approach only provides symmetrical gene-gene interactions without causal relationships and typically yields false positive connections in the network. In contrast, the “extracellular to intracellular model” represents an innovative approach that exploits the spatial information and incorporates the influence of CCC to refine GRN in a spatially correlated manner. NicheNet^28^ and CellCall^29^ predict the communication pathways linking the inside and outside of cells, yet they do not generate spatially constrained GRNs. The recently published CLARIFY utilizes a multi-level graph autoencoder to infer CCC and simultaneously prune GRNs^27^. But it should be noted that CLARIFY has only been applied to a limited number of cells and genes, and it requires an input GRN. On account of the increased sparsity caused by dropouts and lower gene capturing efficiency in SRT data compared to scRNA-seq data, the identification of spatially proximal co-expression patterns across putative cells or spots/bins becomes imperative for predicting statistically significant gene regulation for each cell type or state. It is reasonable to assume that cells exhibiting similar intracellular gene regulation aggerate into spatially constrained groups to perform specific functions in most scenarios. Therefore, one would expect that cells in close spatial proximity would exhibit more similar GRNs under the same intercellular stimulation, while more distant cells would exhibit more distinct GRNs in response to different intercellular signals. Remarkably, none of the existing GRN analysis frameworks for SRT data, to the best of our knowledge, incorporate cis-regulatory motif analysis and spatial gene expression patterns to accurately predict the complete intracellular regulatory paths^30^.

Based on this assumption, we propose the pipeline, SpaGRN, for the inference of spatially-aware intracellular receptor-TF-target network for these locally-distributed scenarios. Considering the continuous increase in the numbers of captured cells/bins and gene as the data size grows, we have employed a proximity-graph-based statistical model with high computational efficiency to investigate potential gene-gene interactions within cells. The spatial proximity and the TF cis-regulatory relationship are jointly used to eliminate false positive connections between TF and targets. In parallel, we identify potential receptors that might be specifically activated by extracellular signals based on the spatially aware co-expression graph, and connect them to intracellular TFs and targets to accomplish the cellular regulation. To evaluate the performance of our pipeline, we have conducted comprehensive benchmarking using simulated SRT data. Our results demonstrate that SpaGRN outperforms several state-of-the-art methods in terms of key evaluation metrics, including the area under the precision-recall curve (AUPRC) and the area under the receiver operating characteristic curve (AUROC). Additionally, our algorithm exhibits superior performance in capturing regulon spatial heterogeneity and offers high computational efficiency. When applied to well-characterized mouse brain datasets, our algorithm recapitulates well-known cell fate regulations governed by TFs, and identifies cell-type-specific regulons with distinct spatial distributions. Moreover, we further interpret regional cell-type-specific regulon configurations and dynamics using a high-resolution 3D *Drosophila* spatiotemporal transcriptomics database, thus enabling mechanistic insights into the cellular and molecular regulation of morphogenesis in the fruit fly.

Furthermore, we have built an interactive 3D GRN atlas database for SRT data (http://www.bgiocean.com/SpaGRN/) rendered by VT3D^31^, that covers different SRT datasets. The availability of this atlas database contributes to advancing related studies, including investigations on embryogenesis and organogenesis processes.

### Design

Briefly, the design of the SpaGRN pipeline, aimed at inferring spatially constrained GRNs from SRT data, encompasses a series of well-defined steps (Figure 1). To begin with, a bipartite spatially-aware gene co-expression network is constructed using the gene coordinated co-occurrence patterns derived from SRT data. This network captures the spatial relationships between genes within spatially proximate cells and serves as the foundation for subsequent analysis. Next, SpaGRN proceeds to infer potential regulatory modules centered around all available TFs. The resulting candidate regulatory modules are subsequently pruned following an approach that is similar to a previously reported method^14^, incorporating promoter region and TF-binding site information. This pruning process eliminates non-significant modules and indirect targets in each significant module. SpaGRN then identifies receptor genes from the co-occurrence graph referring to an established L-R database, and construct final directional receptor-TF-target paths for each TF. The inclusion of receptor genes enriches the biological relevance of the inferred GRNs, accounting for the impact of extracellular signaling on cellular regulation. Finally, cellular receptor-TF-target activities of each regulon are computed in individual cells or spatial bins. This measure can be either enriched to detect cell-identity-specific regulon modules or used for cell clustering to resolve the problem of sparsity for SRT data, which enables the discovery of spatially coherent cell populations and facilitates the interpretation of cellular heterogeneity within the tissue or organ. In summary, by addressing these challenges mentioned earlier and leveraging the power of spatial expression, cell proximity, and extracellular receptor activation, SpaGRN emerges as a robust and effective tool for accurately predicting explicit intracellular regulatory pathways from SRT data.

**Figure 1.**
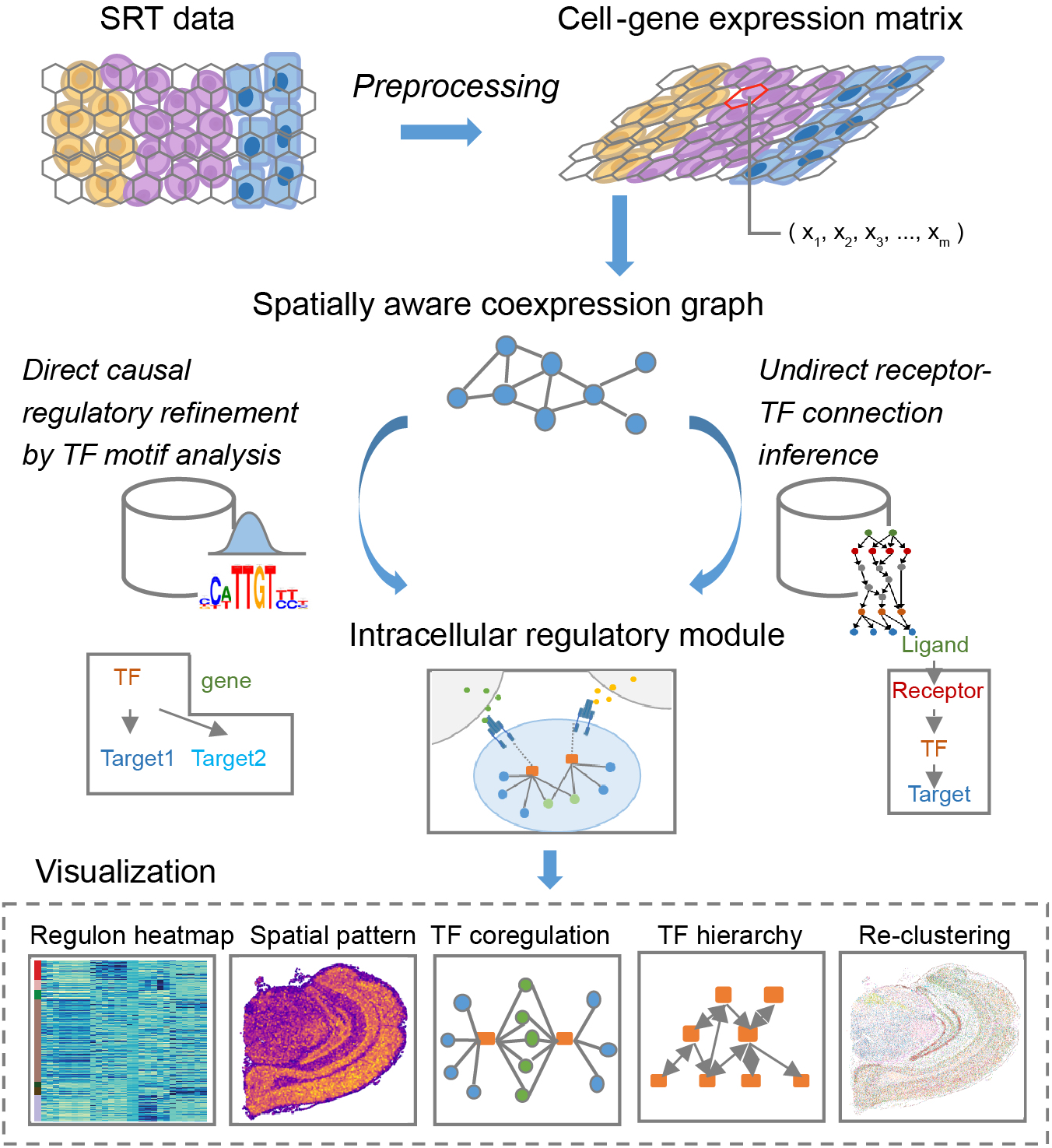
Overview of SpaGRN workflow. First, a bipartite spatially-aware gene coexpression graph is constructed using the preprocessed SRT dataset. Next, SpaGRN infers refined regulatory modules centered around all available TFs, incorporating promoter region and TF-binding site information. In the meantime, receptor genes were identified from the co-expression graph referring to an established L-R database. Finally, SpaGRN constructs a directional receptor-TF-target paths for each TF.

## RESULTS

### SpaGRN identifies spatially resolved regulons

To quantitatively assess the performance on GRN inference of SpaGRN, we generated simulated data using scMultiSim^32^ and applied metrics defined by BEELINE such as AUPRC. These *in silico* datasets included single-cell expressions and locations of TFs and associated targets that were mediated by specific L-R pairs, which served as the ground truth for benchmarking (Figure 2A). We compared SpaGRN against three state-of-the-art GRN inferring methods, namely GRNBoost2, GENIE3, Hotspot. AUPRC and AUROC values were computed as standard classifier scores for a robust comparison. Given that GRNs are typically sparse, the inference problem for these networks exhibits considerable imbalance. It is therefore more appropriate to consider AUPRC as the primary performance metric, rather than AUROC^2,33^ (Figure S1). SpaGRN demonstrated significant performance advantages in accurately inferring TF-target regulations on the simulated data. SpaGRN accurately reconstructed cell cluster-specific GRNs, which exhibited a distribution that correlated with the spatial distribution of corresponding cell clusters (Figures 2B and 2C). To evaluate their stability, we simulated 10 datasets with random differences in technical variations including library preparation noise and batch effects, as well as cell identities involving populations, locations, GRNs, and receptors introduced by the scMultiSim setting. Each algorithm was executed on the 10 replicates, and AUPRC ratios (AUPRC values normalized by that of a random predictor) were calculated. SpaGRN outperformed existing methods across multiple simulated datasets. Specifically, it achieved improvements 41.6%, 31.8% and 48.2% in mean AUPRC ratios compared to GRNBoost2, GENIE3 and Hotspot, respectively, for each of the five individual spatial cell clusters (Figure 2D).

**Figure 2.**
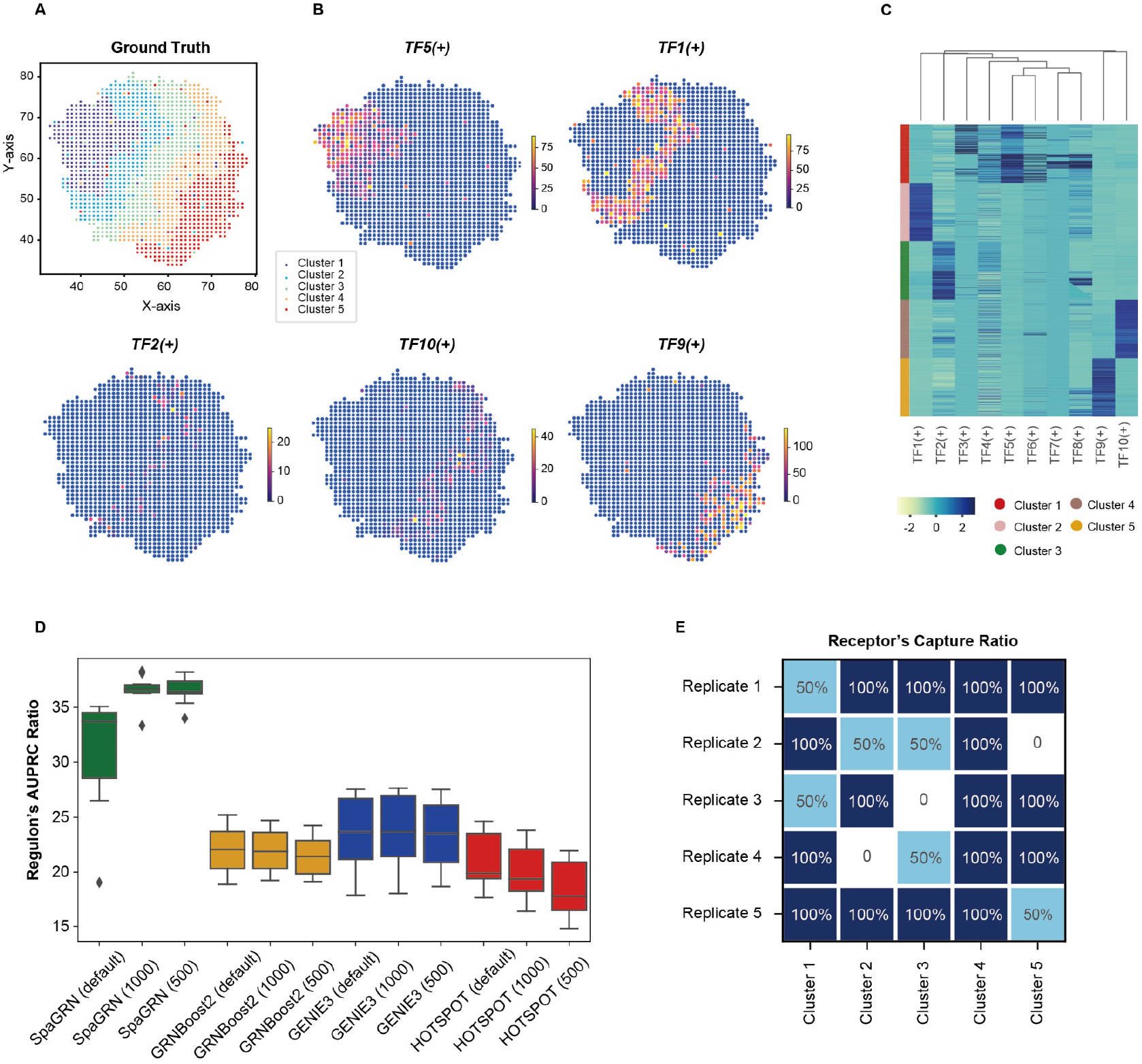
SpaGRN identifies superior spatially resolved regulons with potential intercellular receptors. (A) The spatial distribution of 1500 cells across five cell clusters as groud truth for benchmarking. (B) Detected spatially specific regulons across five cell clusters. (C) Heatmap of functional regulon modules enriched for specific spatial cell clusters. (D) Box-plots depicturing the AUPRC ratio of SpaGRN and three other state-of-the-art GRN inference tools on datasets of different total cell numbers. Each box was generated across ten simulated datasets sampled with scMultiSim. (E) SpaGRN’s capture ratio of ground truth spatial cluster-specific receptors in the five simulated datasets.

We also sought to test the robustness of inferred GRNs obtained through SpaGRN by varying the number of cells to 1500, 1000, and 500 (Figure 2D). The resulting box plots of AUPRC ratios exhibited distinctive dependencies on cell number among the four methods. Nevertheless, SpaGRN demonstrated its superior robustness across varying cell numbers and consistently outperformed the other algorithms across all three sets of ten synthetic datasets investigated.

### SpaGRN investigates intercellular influences on intracellular regulation

Furthermore, we acknowledged that SpaGRN proves to be a valuable tool for not only identifying spatially resolved GRNs but also for elucidating intricate intercellular influences. To evaluate the performance of SpaGRN in capturing relevant receptors to intracellular regulators within the same cell, we utilized scMultiSim simulated datasets that provided a cell-type level CCC ground truth. To assess the stability of capturing receptor genes, we conducted five iterations of dataset simulation, each iteration containing two predefined receptors for every cell cluster for benchmarking purposes. The analysis of the results demonstrated that SpaGRN successfully detected an average of 90%, 60%, 70%, 70%, and 90% of the receptors for the respective five datasets (Figure 2E). This finding highlighted the efficacy of SpaGRN in predicting potential intercellular influences on the intracellular regulons. It is noteworthy that other competing GRN inference methods did not provide CCC information, thus rendering them unable to be compared in this particular aspect.

### SpaGRN unveils spatially resolved regulatory mechanisms underlying brain development

The mouse brain, a well-studied and complex model organ, consists of diverse cell types with distinct gene expression profiles. It provides a unique opportunity to unravel the fundamental cellular and molecular mechanisms governing human brain function, model brain development, and advance neuroscience research. By leveraging SRT data to investigate the spatial organization of gene expression, researchers can decipher networks of TFs, signaling molecules, and other regulatory elements that orchestrate gene expression patterns, contributing to the regional specialization and functional organization of the brain. In this study, we applied SpaGRN to analyze a mouse coronal hemibrain dataset generated using Stereo-seq, aiming to assess its efficacy in detecting spatially resolved regulons.

The cells were clustered and annotated according to differential expression and anatomical structure in the study conducted by Chen et al.^20^ (Figure 3A). Utilizing SpaGRN, we identified highly expressed regions of TFs and associated targets that closely corresponded to the annotated brain regions (Figures 3B and S2). To investigate the regionalization and functional roles of each regulon, we visualized the expression patterns of TFs and target genes, and performed gene ontology (GO) enrichment analysis, with a particular focus on constituent cell types such as excitatory glutamatergic neuron (EX), dopaminergic neuron (DA), and meninge cell types. Notably, the predicted isocortex-specific regulon *Bcl11a*(+) was already known to be present in V1 radial glia and excitatory neurons of the cortical area (Figure S3A). The identified GO terms were directly related to neuron projection morphogenesis, pathways of neurodegeneration, and Alzheimer’s disease, thereby confirming the pivotal role played by *Bcl11a*(+) in regulating neuron progenitor cell proliferation, differentiation, and functional homeostasis^34^ (Figure S3B). Additionally, our findings suggested that *Bcl11a*(+) might maintain its function through the modulation of chemical synaptic transmission, regulation of trans-synaptic signaling, and regulation of calcium ion-dependent exocytosis (Figure S3B). Furthermore, the regulon *Tcf4*(+) exhibited specific expression in the isocortex and hippocampal formation, in line with previous studies that have identified *Tcf4* as a significant marker of glutamatergic and GABAergic neurons in these regions^35^ (Figure S3C). Moreover, further GO analysis revealed a significant correlation between these genes and key processes involved in synaptic signaling and development, providing compelling evidence for their substantial influence on cognitive processes such as perception, attention, learning, and memory^35^ (Figure S3D). We observed the presence of another notable regulon, *Egr3*(+), in both the isocortex and hippocampal formation (Figure 3C), which aligned with previous studies reporting its involvement in excitatory neurotransmission and its ability to modulate neuronal behavior over time (Figure 3D). This regulon was distributed in a distinct Pallium glutamatergic division consisting of the isocortex, hippocampal formation, olfactory areas, and cortical subplate^36,37^. In contrast, DA neuron-specific *Pbx3*(+) exhibited a predominant enrichment in the region of behavioral state related midbrain, which agreed with the reported presence in substantia nigra in midbrain^38^ (Figure 3E). The corresponding GO terms underscored the significance of *Pbx3*(+) in midbrain dopaminergic neuron development, specification and survival, as well as its implication in the development of neurodegenerative diseases^39^ (Figure 3F). Importantly, when compared to the state-of-the-art GRN inference algorithm pySCENIC that has been extensively applied in mouse brain research^20,40^, SpaGRN’s identification was more correlated to real regionalized functions and meanwhile the differentially expressed areas revealed by SpaGRN were more clearly discernible according to the ISH image of TFs obtained from the mouse brain map of the Allen brain atlas (https://mouse.brain-map.org/static/atlas)^41^.

**Figure 3.**
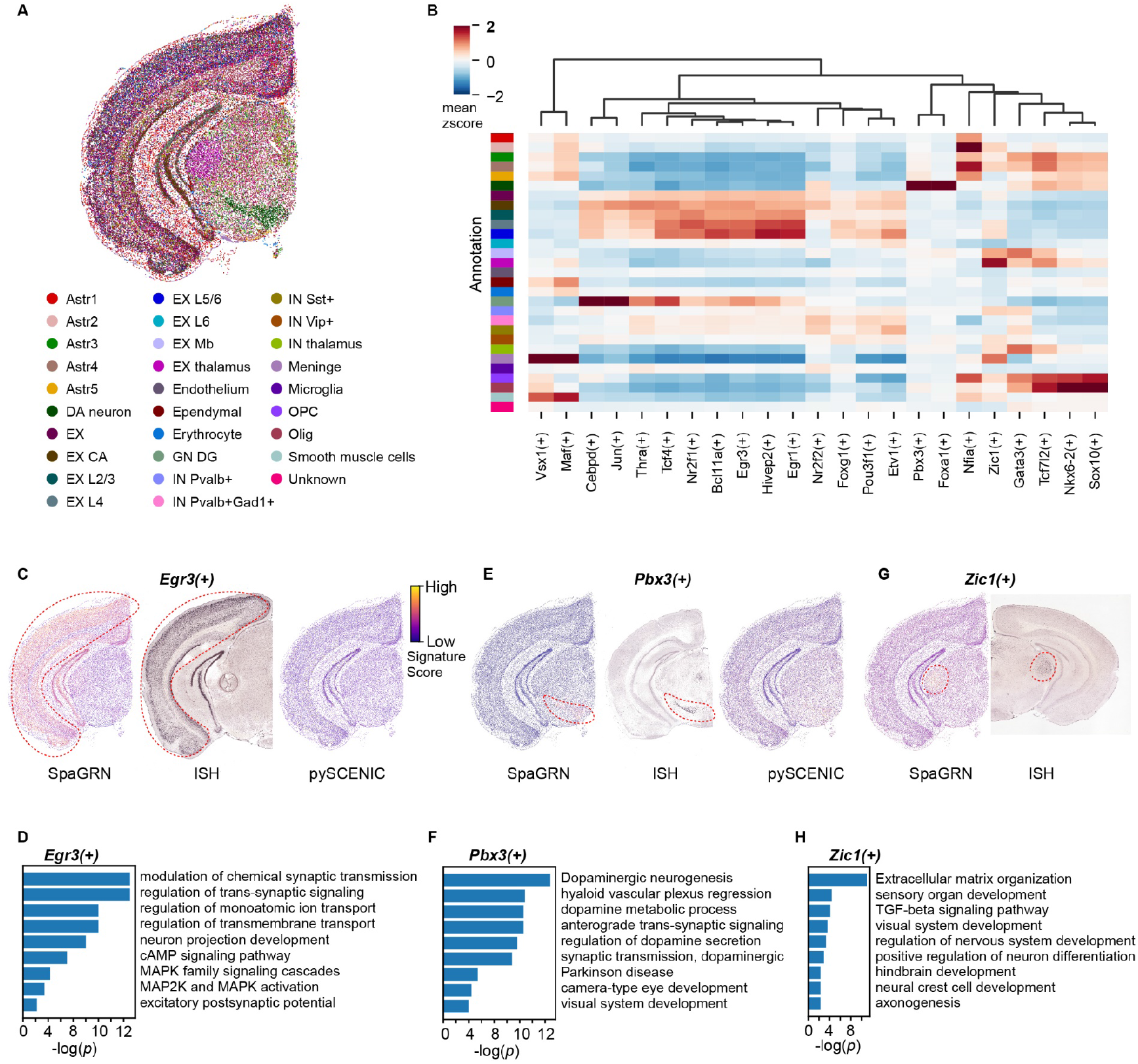
SpaGRN discovers credible and novel regulons with distinct spatial patterns in mouse brain data. (A) Adult mouse hemisphere brain data colored by annotation. (B) Heatmap of functional regulon modules enriched for specific cell types. (C, E) Spatial representation of regulons *Egr3*(+) and *Pbx3*(+) using SpaGRN and pySCENIC. The spatial distribution is verified by the corresponding ISH from the mouse brain map of the Allen brain atlas. (D, F) GO enrichment analyses of *Egr3*(+) and *Pbx3*(+). (G) Spatial representation of regulon *Zic1*(+) uniquely detected by SpaGRN and the corresponding ISH image for validation. (H) GO enrichment analysis of *Zic1*(+).

In the spatial context, cellular heterogeneity is determined not only by the intracellular regulatory network but also by the extracellular microenvironment, which collaboratively accomplish diverse biological functions^42,43^. However, existing computational methods for modeling both interactions simultaneously are still underdeveloped^44^. SpaGRN exhibited an exceptional capability to precisely identify potential receptors that might contribute to regulon activity, providing unique insights into intracellular regulation mediated by biochemical signals and intercellular crosstalk across multiple dimensions. Within several EX cell clusters, our findings suggested that intercellular communication through specific receptors potentially regulated the activity of *Tcf4*(+). Among them, previous studies have reported the co-expression of *Cntn5*^45^, *Stx1a*^46^, *Scn2a*^47^, *Thy1*^45^, and *Scn1b*^48^ with *Tcf4*(+). Moreover, SpaGRN predicted the involvement of previously unreported receptors, *Il31ra* and *Gp1bb*. Furthermore, the regulon *Bcl11a*(+) associated with neuron proliferation and differentiation, was inferred to be functionally linked to known co-expressed receptors, including *Nptxr*^45^, *Negr1*^49^, *Pde1a*^49^, *Thy1*^45^, and *Lrrtm4*^50^. Additionally, *Scn1b*, *Il31ra*, *Mpzl2*, and *Asgr1* were identified by SpaGRN as potential mediators of *Bcl11a*(+) activity, providing novel insights into their regulatory roles that have not been previously reported. Within the EX cells in isocortex and hippocampal formation, the known receptors *Adcy1*^51^ and *Camk2a*^51^ were identified as key receivers of extracellular signaling, influencing the intracellular activity of *Egr3*(+). This effect could also be attributed to the co-expression of *Tspan13*, *Stx1a*, *Atp6ap2*, *Opcml*, *Scn1b*, *Pde1a*, *Lingo1*, *Lrrtm4*, *Camk2a*, *Scn2a*, *Gp1bb*, and *Sv2b*. Moreover, the midbrain-specific *Pbx3*(+) was associated with *Chrna4*^45^ and *Ret*^45^, with the additional discovery of new receptor *Chrnb3*. SpaGRN provided researchers with direct signaling pathways that bridged inter- and intracellular regulation, connecting selected receptors, TFs, and targets, thereby streamlining the research process and minimizing labor-intensive experimentation.

SpaGRN presents a comprehensive and robust solution that surpasses existing GRN inference algorithms such as pySCENIC, for analyzing complex SRT data. In the mouse brain dataset, SpaGRN revealed additional spatially resolved regulons, including *Zic1*(+), *Gata3*(+), *Hivep2*(+), and *Vsx1*(+), which were not identified by pySCENIC. Notably, the EX thalamus and Meninge-enriched *Zic1*(+) emerged as a thalamus-specific regulon, known for its significant role in maintaining neural precursor cells in an undifferentiated state^52^ (Figure 3G). Furthermore, GO enrichment analysis confirmed its involvement in the development of sensory organs, the visual system, hindbrain, and neural crest cells (Figure 3H). Importantly, our findings suggested that these functional associations were mechanistically linked to extracellular matrix organization and the TGF-beta signaling pathway (Figure 3H). Additionally, *Zic1*(+) was predicted to be regulated by a set of known co-expressed receptors including *Fgfr2*^53^, *Gjb2*^54^, *Gpr182*^55^, *Thbd*^56^, *Cdh1*^57^, *Ramp3*^58^, *Cd44*^59^, and *Anxa2*^60^, and we also identified previously unreported receptors *Il31ra*, *Mpzl2*, and *Asgr1*. Another regulon, *Hivep2*(+), exhibited a distribution primarily enriched in the frontal cortex and hippocampus^61^ (Figure S3E). The corresponding GO terms aligned well with its crucial role in severe cognitive and social impairments, anxiety-like behaviors, hyperactivity, and memory deficits^61^ (Figure S3F). Additionally, the spatial distributions of highly expressed regions of *Gata3*(+) (Figure S3G) and *Vsx1*(+) (Figure S3H) were in accordance with previous studies^46,62^. Their distributions were further supported by corresponding ISH benchmarking images.

In conclusion, SpaGRN offers a rigorous and systematic analysis of inter- and intracellular signaling pathways, yielding invaluable insights into the intricate regulatory mechanisms underlying brain structure and function. By employing the SpaGRN algorithm and elucidating the roles of receptor-mediated regulons, significant advancements have been made in our comprehension of the complex biological processes orchestrating brain development and activity. This comprehensive approach not only enhances our understanding of brain function at cellular and molecular levels but also opens up promising avenues for therapeutic interventions and the treatment of neurological disorders.

### SpaGRN observes spatiotemporal regulatory variations in time-series 3D SRT data

Organismal growth and development involve complex biological processes characterized by variations in cell types and gene expression over time. These temporal variations capture the dynamic molecular changes occurring during development. In this study, we leveraged a previously published 3D square-binned atlas of *Drosophila* embryogenesis^23^ to emphasize dynamic changes in both the spatial and temporal dimensions, encompassing 5 developmental stages that span embryos and larvae.

We first inferred the spatial GRN configurations in the 5 stages and detected 33, 22, 22, 25, and 14 regulons, respectively. Through unsupervised clustering followed by manual annotation, we identified regulons enriched in different cell types (Figures 4A and S4). These regulons including central nervous system (CNS)-specific *hth*(+), midgut-specific *cad*(+), and epidermis-specific *vvl*(+), showed spatial patterns similar to the reconstructed 3D organ mesh models at different developmental time points (Figure 4B). Subsequent ISH examinations also confirmed that *hth*, *cad*, and *vvl* were mainly expressed in the CNS, midgut, and epidermis region, respectively (Figure 4C). By applying SpaGRN to calculate spatiotemporally dependent regulon activities, we identified the dynamic functions of these cell-identity-specific regulons imperative for embryogenesis and organogenesis. In particular, our observations revealed the dynamic migration of the highly expressed region of the *hth*(+) regulon from the neurogenic ectoderm region during early embryonic stages to the head region during larval stages. This spatial shift was consistent with the dynamic regulation of neuron differentiation and morphogenesis, as indicated by GO enrichment analysis^63,64^ (Figure 4D). Another regulon, *cad*(+) successively experienced expansion and contraction during midgut development. Although the GO enrichment analysis failed due to the limited number of captured targets, it is worth noting that previous studies have indicated the involvement of this TF in transmembrane transport and chemical homeostasis, implying a potential role in digestion and absorption^65^. In contrast, the regulon activity of the epidermis-specific *vvl*(+) remained in the ectoderm region throughout development. Its enriched GO terms were primarily related to epithelial morphogenesis as previously described^66^ (Figure 4E). Compared to the GRN results from the original article^23^, SpaGRN obtained regulons with more evident and precise spatial heterogeneity for the same TF with the benefit of its spatial refinement capability (Figure S5).

**Figure 4.**
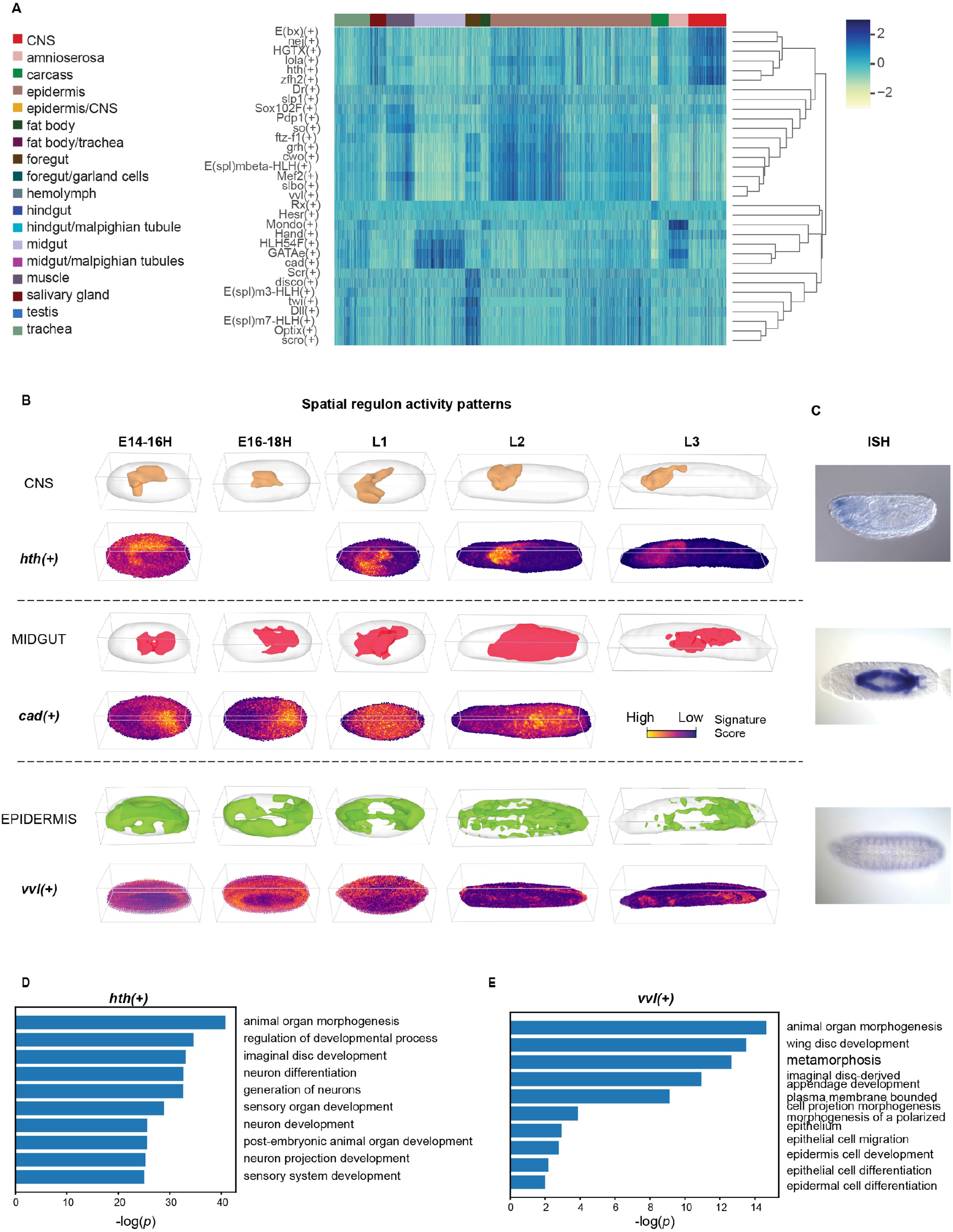
SpaGRN identifieds 3D spatiotemporal patterns of regulons for developing *Drosophila*. (A) Heatmap of functional regulon modules enriched for specific cell types. Left color bars refer to different cell types. (B) Cell-identity-specific regulons during different developmental stages. The typical spatial regulon (bottom) for different cell types shows a highly similar pattern in 3D to the morphological mesh model of the corresponding organ or tissue (top). (C) ISH experiments validating the spatial distribution of the TF for each specific regulon. (D) GO term enrichment analysis of representative regulons in (B).

Moreover, our investigation delves into the co-regulatory connections among different specific regulons enriched in the same tissue. For instance, we inspected the crucial cooperative role of two CNS-specific regulons, *hth*(+) and *exd*(+) in neuron differentiation. These two regulons were connected through shared target genes, and could be collectively stimulated by the receptor *msn*, or independently by *LRP1*, *Egfr*, *ATP6AP2*, *mag*, *robo3*, *Lamp1*, *Alk*, in response to extracellular signaling (Figure 5A). The correlation between *hth*(+) and *exd*(+) has been reported, with *hth* responsible for the *exd* retention in the nucleus and forming part of the functional Exd/Hox complex^67^. Double immunostaining experiments further verified the co-expression of these two regulators in olfactory projection neurons and their progenitors^68^. GO term analysis of shared targets of these regulons also indicated their vital roles in neuron differentiation, aligning with the reported function of encoding a MEIS family protein that regulated the subcellular localization of the homeotic protein cofactor Extradenticle, involved in multiple aspects of embryonic and adult fly development^63,64^ (Figure 5B). The mutation of *hth* or *exd* has been shown to result in defects in olfactory projection neurons^68^. On the other hand, the identification of unique targets for each regulon implied their subtle distinctions in regulatory functions (Figure 5B).

**Figure 5.**
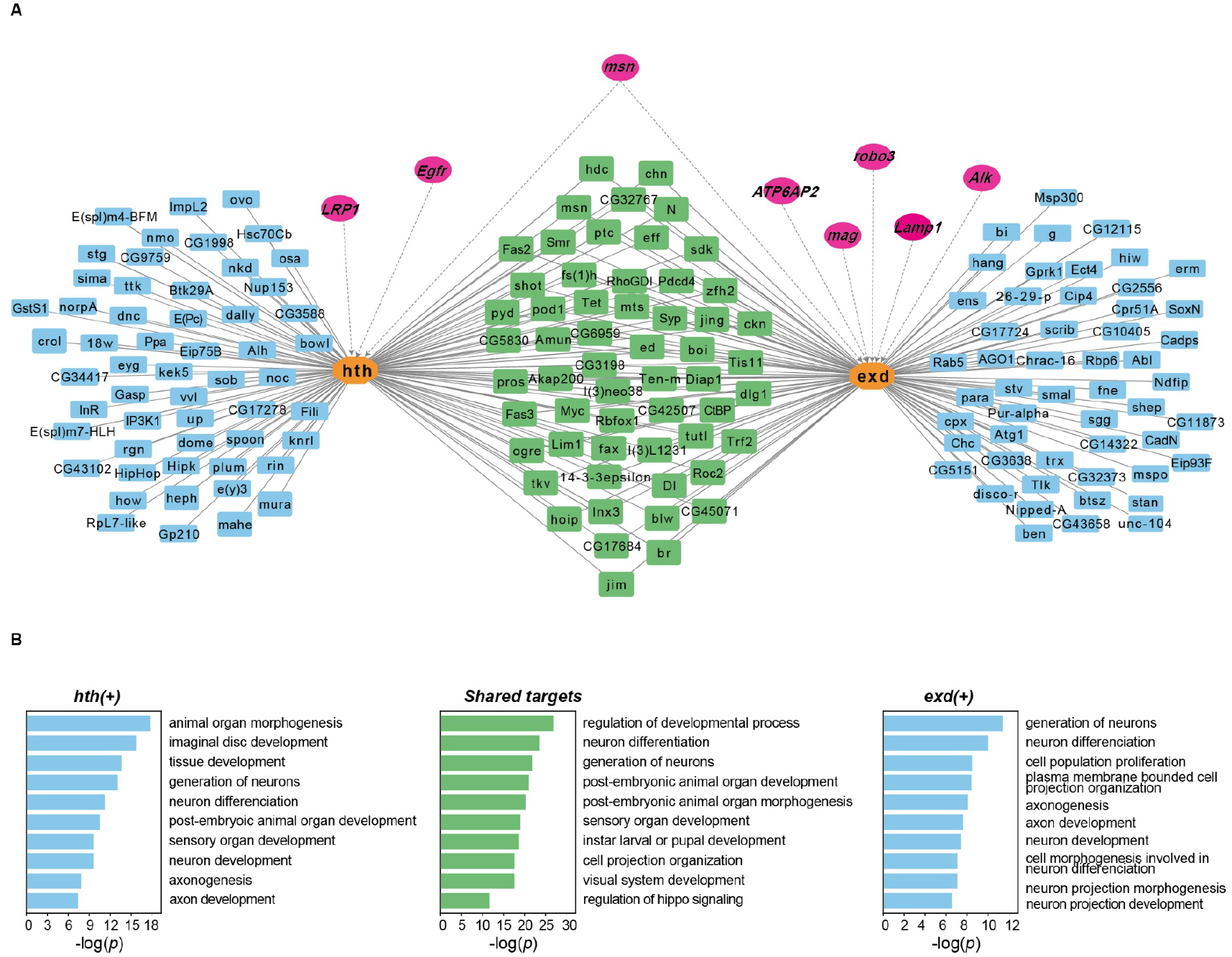
SpaGRN unveils an intra- and inter-cellular co-regulation network in CNS. (A) Connection network of two representative spatially specific regulons enriched in CNS. The detected influential receptor genes are shown in magenta color and points to the TFs with dashed arrows. (B) GO terms of shared and exclusive target genes in *hth*(+) and *exd*(+) regulons.

Therefore, SpaGRN facilitates the investigation of specific receptors that transmit signals from CNS’s niche, which promote or inhibit the activity of specific TFs inside CNS cells through specific signaling pathways. These TFs ultimately influence the expression of downstream targets, collectively contributing to the cellular functions relevant to organogenesis.

### Performance

GRN inference from large-scale gene expression data of SRT or scRNA-seq has been computationally demanding. To evaluate the computational performance of SpaGRN, we compared it to two existing algorithms, pySCENIC and Hotspot, using the same mouse hemibrain dataset consisting of 50,140 cells and 25,879 genes (Table 1). The current pySCENIC pipeline, implemented in Python, claimed to have increased the computational efficiency compared to its predecessor by introducing XGBoost and parallelization. However, our results demonstrated that SpaGRN exhibited an average 5- and 22-fold increase in the running speed compared to pySCENIC and Hotspot, respectively, when running on a Linux system equipped with an Intel Core Processor (Broadwell, IBRS) of 40 CPU threads and 256 Gb RAM. The peak memory usages of the three methods were similar, though. We noted that other GRN tools such as CellTrek and SpaceX required significantly longer computation times (more than 2880 CPU hours) when tested on the same dataset. In addition, the performance of SimiC was heavily dependent on user-defined parameters, such as the number of output regulons and target genes. These tools were not included in our benchmarking analysis. We also tested the performance comparisons using the fruit fly data (Table S2).

**Table 1.** Statistics of computational performance on mouse brain data.

| tools | clock time (hour) | peak memory (Gb) |
| --- | --- | --- |
| SpaGRN | 2.2 | 67 |
| pySCENIC | 9.2 | 82 |
| Hotspot | 39 | 68 |

The GRN inferences were performed on a computer cluster with 40 Intel Core Processor (Broadwell, IBRS) CPU threads with 256 Gb of memory installed, and used the Linux operating system.

## DISCUSSION

The burgeoning field of single-cell omics has provided unprecedented opportunities for modeling and predicting the regulators of cellular identities. In this paper, we have introduced SpaGRN as a versatile approach for exploring directional GRNs in SRT data. This pipeline capitalizes on the integration of spatially-aware transcriptomic profiles, TF motif sequences, and L-R-TF linkages to infer cell-identity-specific regulon modules. By leveraging benchmarking analyses and practical applications, we have demonstrated the efficacy of SpaGRN in reconstructing robust guides that elucidate regulatory mechanisms within specific cells or regions, thereby capturing topological patterns inherent in tissue sections. Furthermore, SpaGRN exhibits remarkable flexibility, accommodating mainstream coordinated cell-gene matrices or cell-gene similarity graphs generated by most SRT platforms. Notably, our framework can be extended to incorporate such matrices or graphs produced by computational algorithms that map single cells back to their spatial coordinates through the integration of scRNA-seq and SRT data. Additionally, SpaGRN offers users the convenience of visualizing and exploring spatial GRNs through flexible 2D imaging functions and an interactive 3D online atlas database provided within the framework (Figure S6).

### Limitations

While SpaGRN represents a powerful tool for exploring spatially informed GRNs, it is important to acknowledge its inherent limitations. One such limitation is the potential presence of false-positive or false-negative interactions within the inferred spatial GRN, emphasizing the need for experimental validation to confirm the reliability of the network predictions. Although SpaGRN leverages disparate forms of information such as spatial transcriptomic profiles, TF-target cis-regulatory, and existing receptor-TF linkages, it is crucial to recognize that certain relevant biological interactions that occur in a non-cell-autonomous manner or between distant cells may not be effectively captured by the current framework.

Similar to other computational methods, SpaGRN also heavily relies on the quality and reliability of the input data. Inaccuracies, noise, or biases in SRT profiles can impact the accuracy and robustness of the inferred spatial GRNs. The spatial resolution of SRT data can also pose limitations on the ability to capture fine-grained regulatory interactions. Moreover, the accuracy of SpaGRN’s predictions is contingent upon the availability and comprehensiveness of annotated TF-target and L-R-TF in the two knowledge libraries. Incomplete or biased annotations may result in the overlooking of important regulatory interactions, potentially leading to an incomplete representation of the spatial GRN and associated signaling paths. At last, the performance and generalizability of SpaGRN may vary across different tissues and species. One main reason is the two existing knowledge libraries were pre-trained by model organisms, potentially affecting the applicability and accuracy of SpaGRN in rare organs or in non-model organisms.

Moving forward, we have outlined several directions for future research to enhance the capabilities of SpaGRN. One avenue of investigation involves tracing the propagation of regulatory signals within the spatial GRN. By elucidating the flow of regulatory information such as pseudotime regulatory trajectory tracing^14^ across cells and regions, we aim to gain valuable insights into the spatiotemporal dynamics of gene regulation. Additionally, we plan to conduct *in silico* gene perturbation experiments^16^ within the SpaGRN framework. This simulation-based approach will allow us to investigate the functional consequences of perturbing specific genes or regulatory elements within the inferred GRN, enabling us to predict the impact of gene manipulations on cellular behavior and regulatory outcomes.

## ACKNOWLEDGMENTS

We thank Hanbo Li, Li Deng, Yong Zhang, and Yuxiang Li for the helpful discussion and critical reading of the manuscript. We also thank all the public databases contributing to this work including CNGBdb, STOmicsDB and SODB for data sharing. This work has been supported by the National Natural Science Foundation of China (No. 32100514); the General Program (Key Program, Major Research Plan) of National Natural Science Foundation of China (No. 32170439); the National Natural Science Foundation of China (No. 32300526); and National Key R&D Program of China (No. 2022YFC3400400).

## AUTHOR CONTRIBUTIONS

M.X. and G.F. conceived and directed the study. M.X., G.F., and X.X. supervised the work. Y.L., M.X., X.L., and L.G. developed the SpaGRN method. Y.L., X.L., L.G., and K.H. generated and analyzed the data. L.G. and Y.L. performed database construction. S.F. and X.W. assisted in the data analysis. M.X. drafted the manuscript. Y.L., X.L., L.G., S.F., X.X, and G.F. edited the manuscript.

## STAR METHODS

## RESOURCE AVAILABILITY

### Lead Contact

Further information and requests for the resources and reagents may be directed to and will be fulfilled by the lead contact, Mengyang Xu.

### Materials availability

This study did not generate new unique reagents.

### Data and code availability

● All data used in this study have been previously reported and are publicly available (Table S1). The coronal hemibrain section in an adult mouse is available from STOmicsDB MOSTA database^69^ (https://db.cngb.org/stomics/mosta/download/). The raw sequencing data are stored in CNGB Nucleotide Sequence Archive (CNSA)^70^ of China National GeneBank DataBase (CNGBdb)^71^ under accession number CNP0002646. The 3D developing *Drosophila* embryos and larvae dataset was downloaded from STOmicsDB (https://db.cngb.org/stomics/flysta3d/). The raw sequencing data is available in the CNGB under accession number CNP0002189. Processed data can be interactively explored from our database (https://www.bgiocean.com/SpaGRN). All the related data and TF databases can be directly obtained from our database website.
● All original codes supporting the current study are hosted on GitHub (https://github.com/BGI-Qingdao/SpaGRN).
● Any additional information required to reanalyze the data reported in this paper is available from the lead contact upon request.

## QUANTIFICATION AND STATISTICAL ANALYSIS

### Data design and simulation for benchmarking

We evaluated the GRN inference ability of SpaGRN and existing methods on ten replicates of simulated SRT datasets, each consisting of a single-cell gene expression matrix of 1800 genes and 1500 cells across five clusters, along with the spatial coordinates of cells. We used scMultiSim^32^ to generate these simulated datasets based on a pre-defined ground truth GRN consisting of five regulons with distinct spatial patterns and another five regulons uniformly distributed throughout the spatial area. Furthermore, we assessed the receptor detection ability of SpaGRN using another five simulated SRT datasets generated with scMultiSim. Each of these five datasets involved 120 genes and 1500 cells of five types and was simulated based on a pre-defined ground truth GRN of five spatially specific regulons as well as 20 pre-defined ground truth L-R pairs.

The first ten simulated gene expression datasets were generated using scMultiSim through a two-step process. During the first round of simulation, we employed the ‘layers’ layout and generated coordinates for a total of 1500 cells belonging to five different clusters (300 cells for each cluster) by specifying 20 L-R pairs in the ‘cci’ parameter (Figure 2A). In the second round, we independently simulated five sets of expression data, each consisting of 300 cells and 1800 genes and using one of five pre-defined regulon parameters. These five sets of cells were assigned to the coordinates corresponding to the five clusters that were generated in the previous round of simulation. In this way, these five regulons could be considered as spatially specific. In addition, we generated gene expression data of five spatially non-specific regulons by simulating a dataset of 1500 cells and 1800 genes with five additional regulon parameters and assigning the 1500 cells randomly to these 1500 coordinates. The resulting dataset comprised a total of 1500 cells, evenly distributed among five distinct cell types that were spatially separated from one another. The 1800 genes include a combination of five cell-type specific regulons and five generally existing regulons, as well as some non-regulon-related noise.

The other five simulated datasets were also generated with scMultiSim. We utilized the same 20 L-R pairs and five different regulon parameters. The resulting datasets each consisted of 1500 cells distributed across five spatial clusters and a total of 120 genes.

### Real data collection and preprocessing

We downloaded the 2D SRT dataset of the coronal hemibrain section in an adult mouse. This dataset was generated by Stereo-seq and has been extensively studied as a reliable benchmark for evaluating the pipeline’s performance. The data captured mRNA expression from 25,879 genes in 50,140 putative single cells with spatial locations.

Furthermore, we applied SpaGRN to systematically analyze the intracellular regulation across the fruit fly development using the 3D developing *Drosophila* embryos and larvae dataset. Since no cell segmentation information was available, we used 15,295 bin20 (20 spots × 20 spots) with 13,668 genes, 14,634 bin20 with 12,850 genes, 17,787 bin20 with 13,083 genes, 64,658 bin20 with 14,270 genes, and 43,310 bin50 with 16,326 genes for E14-E16, E16-E18, L1, L2, and L3 samples, respectively.

### Spatially-aware gene co-expression graph construction

The proposed SpaGRN starts with the construction of a spatially-aware gene co-expression graph. Traditional computational co-expression methods, such as correlation-based statistics, Bayesian networks, and machine learning approaches, are commonly used for scRNA-seq data and can be straightforwardly applied to SRT data. But the incorporation of spatial co-occurrence and co-distribution relationships remains largely underdeveloped. To address this, we leverage the power of a graph structure to make full use of spatial information. The similarity graph has been previously demonstrated to be effective in detecting gene modules in multimodal settings for SRT data^24^. In this work, we utilize the K-nearest-neighbors (KNN) graph as our similarity graph. Each cell is represented as a node, and we connect each cell to its K nearest cells based on the Euclidean distance.

Using this spatial-proximity graph, SpaGRN calculates spatial autocorrelation to evaluate whether the expression of a given gene in a cell can be predicted by its neighboring cells in the graph using the statistic

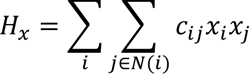

where *x_i_* is the standardized expression of the gene in cell *i*, *N*(*i*) is the K nearest neighbors of cell *i*, and *c*_*ij*_ is a non-negative weight determined by the distance between cell *i* and *j*. The calculated value is dependent on the variation in gene expression patterns across different regions in the spatial-proximity graph. The dependence of each gene on the graph structure can be statistically tested with a parametric null model assuming that the gene expression values in each cell are independently drawn from a certain distribution, without any spatial autocorrelation. Here, we employ the negative binomial distribution as the selected null model and compute the Z scores of *H*_*x*_, from which we can determine the significance of each gene.

We retain only the genes with significant spatial patterns based on the *H_x_* statistic, and divide them into TF and non-TF genes referring to a built-in or user-defined whitelist. Subsequently, we build a bipartite co-expression graph connecting every TF with all the non-TF genes. The weight assigned to each TF-target edge is determined by calculating a local co-expression score, which is defined by the following formula:

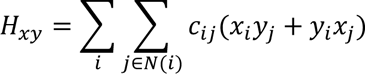

where *x_i_* and *y*_*i*_ are the standardized expressions for gene *x* and *y* in cell *i*. Then for every TF in the co-expression graph, we apply multiple selection strategies, such as TF’s top *N* targets or top *n*th percentile of targets, to construct multiple regulatory modules centered around it.

### Direct causal regulatory refinement by TF motif analysis

Next, we filter the previously constructed regulatory modules and prune the unauthenticated connections from the retained regulatory modules. Given that the TF-centered regulatory modules are generated through spatial proximity screening, they may still contain indirect and potentially misleading targets. Since multimodal data are not always be readily available for deriving the true TF-gene interactions, we integrate gene expression and prior knowledge through the cis-regulatory motif enrichment analysis for direct regulatory inference. We perform cisTarget^72^ to sort and rank all target genes of each potential regulatory module. Using the hidden Markov model and positional weight matrices, the motif enrichment score of a module is calculated based on the area under the ranking curve (AUC) value. Only modules whose target genes show significant higher motif enrichment score are retained. Meanwhile, we also leverage the cisTarget rank to prune the target genes in every retained module. To enhance user convenience, we have downloaded three pre-computed motif databases for three organisms, *Homo sapiens*, *Mus Musculus*, and *Drosophila* melanogaster (https://resources.aertslab.org/cistarget/)^72^ and included them in our pipeline. Given a gene set, this analysis aims to further refine the topological structure of the inferred GRN such that the edges represent direct interactions corresponding to activation or inhibition from the TF to its target genes.

### Undirect receptor-TF connection inference

The intracellular GRN is not isolated from the cellular microenvironment. Potential intercellular communications can be inferred with any CCC tools for SRT data. To investigate the CCC effect from neighboring cells that may impact the TF-centered GRN, we retrieve receptors from the co-expressed genes of each TF that were excluded during the motif analysis. Subsequently, we link these receptors to their corresponding TFs within the same cells, revealing indirect connections between receptors and TFs. In this study, we utilize the NicheNet V2.0 L-R network database for *Homo sapiens* and *Mus Musculus* (https://zenodo.org/record/7074291) to determine the receptors, which integrates 14, 23, and 20 database sources for L-R interactions, signaling paths, and TF regulatory relationships, respectively. In total, it includes 1,430 L-R pairs, 18,647 signaling linkages, and 25,108 regulatory interactions.

### ISR score at single-cell resolution

To evaluate the receptor-TF-target module as a whole at the single-cell or binned resolution, we define and calculate its ISR score in every individual cell. This scoring approach is inspired by the recovery-framework-based AUCell algorithm implemented in SCENIC^14^. It is robust against dropouts and noise frequently observed in SRT data through measuring the enrichment of each receptor-TF-target signature in a ranking-based manner, rather than using the original gene expression profiles.

First, we rank all the genes in a given cell in descending order based on their expression levels. Genes with the same expression value are shuffled. Let *S*_*k*_ = {*g*_1_, *g*_2_,…*g*_*nk*_} be the gene set of the *k*_*th*_ receptor-TF-target signature, where *n*_*k*_ is the number of genes in the *k*_*th*_ signature. Let *rank*(*g*_*i*_) = *r*_*i*_ be the rank of gene *i* in a cell. For a pre-defined critical threshold *r_thre_*, let 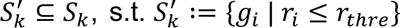. Then, the ISR score of the signature *k* is defined as

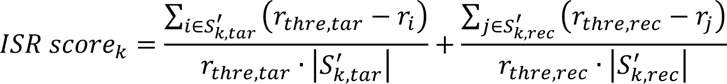

where *tar* refers to all the target genes while *rec* means all the receptor genes. This ranking-based gene signature scoring represents a dimensionless quantity, which is robust to the data processing preferences such as the choice of normalization procedure. More importantly, it extended the traditional concept of TF-target GRN to encompass both targets and receptors that are co-expressed with TFs inside the same cell, building up a more comprehensive picture of the intracellular signaling regulation network.

The resulting spatial cellular ISR matrix has reduced dimensions but retains most informative regulatory profiles, and can be used for significant regulatory subnetwork identification across different cell types and functional regions. The cell-type- or cell-state-specific regulons are finally obtained via enrichment analysis using the hierarchical clustering algorithm.

### Static and interactive visualizations

SpaGRN accepts two commonly used HDF5 file formats, loom and H5AD, for keeping and managing large and complex data. Written in Python language, SpaGRN can effectively handle large SRT datasets with its multiprocessing capabilities, making it well-suited for big data applications. In addition, SpaGRN offers diverse static and interactive visualizations ascribable to a variety of libraries including Matplotlib^73^, Seaborn^74^, and Plotly^75^. It is also convenient to integrate SpaGRN with other SRT or scRNA-seq data analysis toolkits such as Scanpy and Squidpy^76^. We note that SpaGRN has been added to the latest version of Stereopy, which is a fundamental and comprehensive SRT analysis toolkit (https://github.com/STOmics/Stereopy).

### Benchmarking methods for regulon reconstruction

We applied the BEELINE^2^ framework to benchmark GRN results using different inference methods including Hotspot, GENIE3 and GRNBoost2 on the simulated datasets. We ranked every possible TF-target interaction edge in the output GRN and converted them to a ranked list. According to the ground truth GRN and CCC, the AUPRC and AUROC values were computed as metrics from the SpaGRN’s local autocorrelation coefficients, GENIE3’s regulatory weights, GRNBoost2’s importance scores, and Hotspot’s autocorrelation scores.

