## supplementary information for "SpaGRN: investigating spatially informed regulatory paths for spatially resolved transcriptomics data"

### *Supplementary Material*

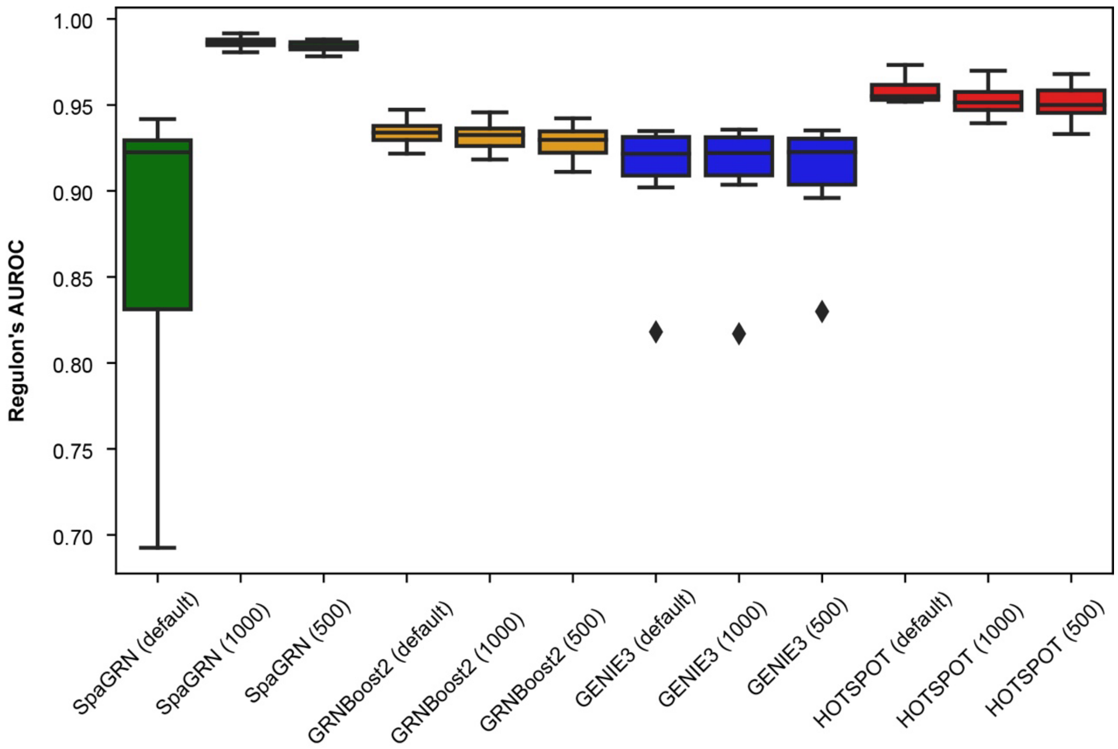


**Figure S1. AUROC values of different GRN inference tools, related to Figure 2.** Box-plots depicturing the AUROC values of SpaGRN and three other state-of-the-art GRN inference tools on datasets of different total cell numbers. Each box was generated across ten simulated datasets sampled with scMultiSim.

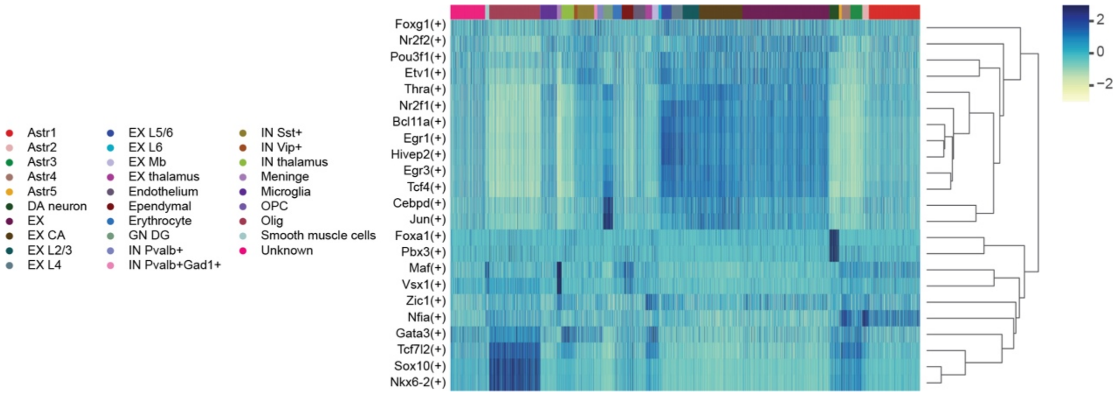


**Figure S2. Heatmap of cellular regulon activity scores enriched in specific cell types for adult mouse coronal hemibrain, related to Figure 3.**

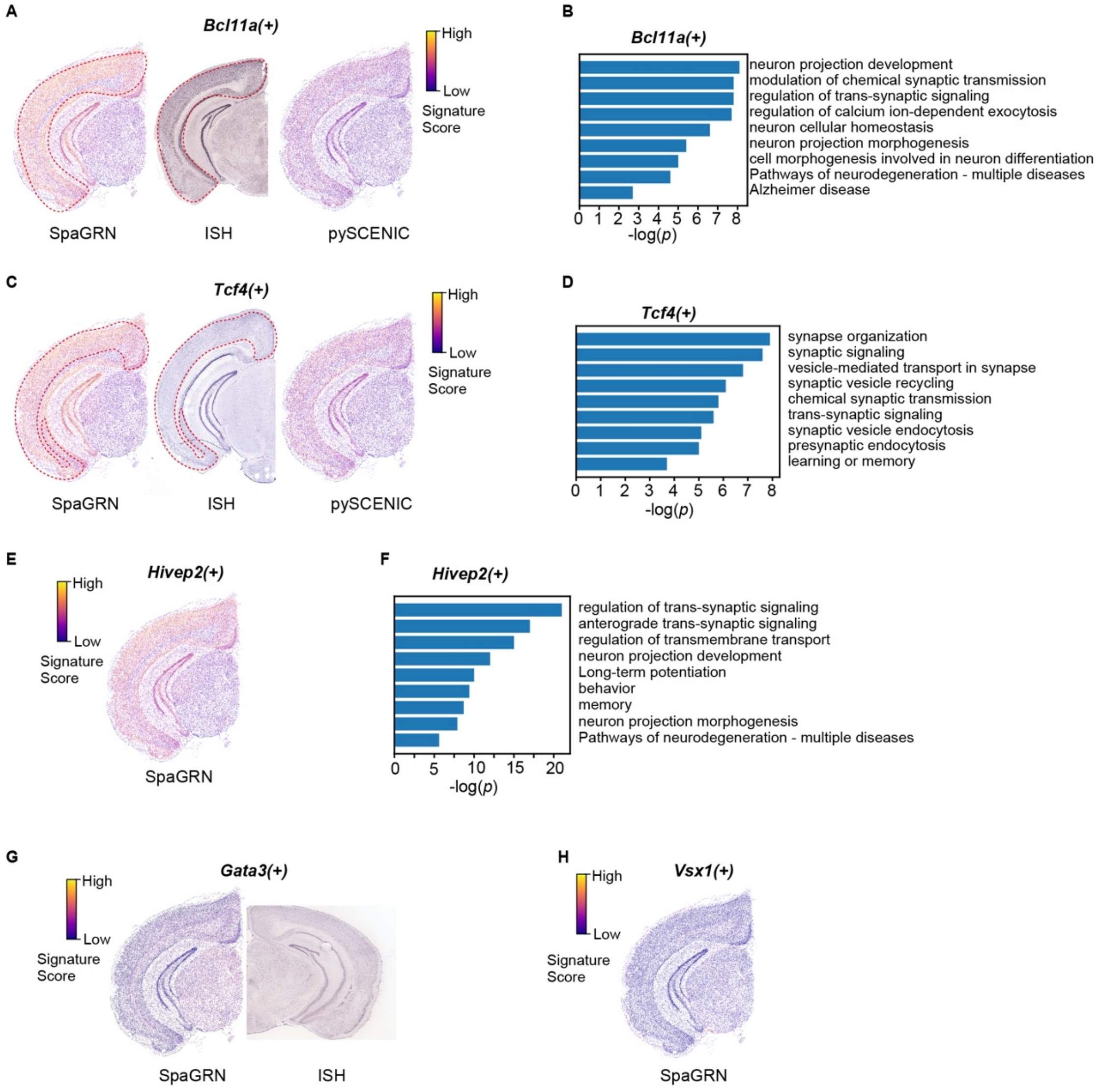


**Figure S3. SpaGRN discovers credible and novel regulons with distinct spatial patterns in mouse brain data, related to Figure 3.**

(A, C) Spatial representation of regulons *Bcl11a*(+) and *Tcf4*(+) using SpaGRN and pySCENIC. The spatial distribution is verified by the corresponding ISH from the mouse brain map of the Allen brain atlas.

(B, D) GO enrichment analyses of *Bcl11a* (+) and *Tcf4*(+).

(E,G,H) Spatial representation of regulons *Hivep2*(+), *Gata3*(+) and *Vsx1*(+) uniquely detected by SpaGRN and the corresponding ISH image for validation.

(F) GO enrichment analysis of *Hivep2*(+).

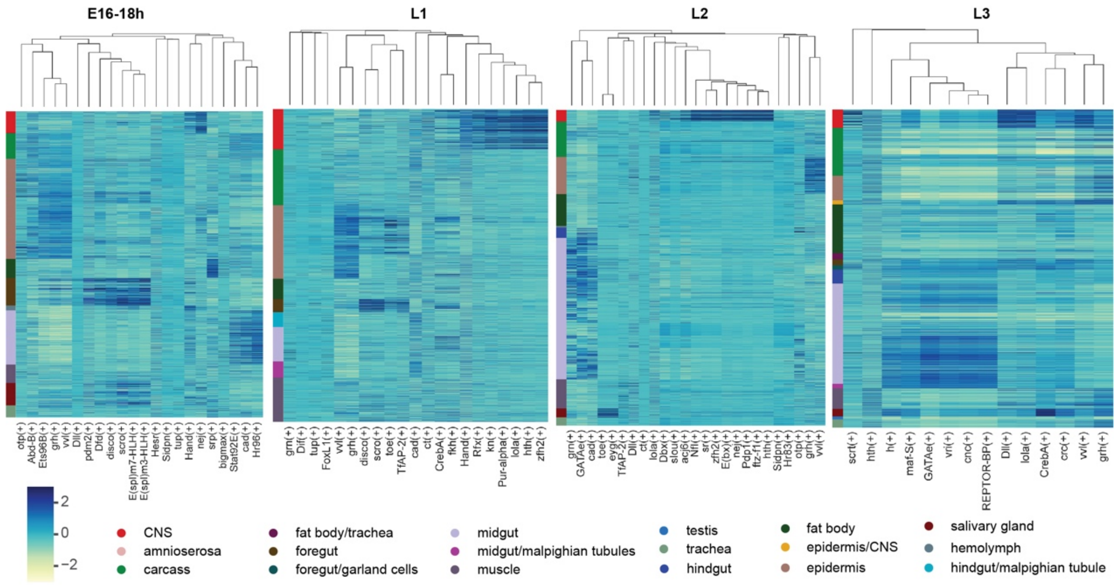


**Figure S4. Heatmaps of cellular activity scores for cell-type-specific regulons from developing *Drosophila* data, related to Figure 4.**

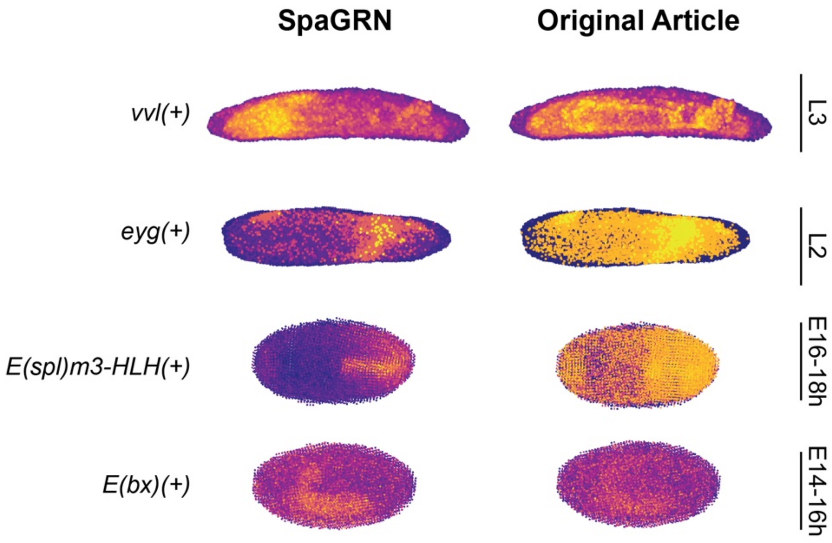


**Figure S5. Comparison of regulons identified by SpaGRN and original article (pySCENIC) at different *Drosophila* developmental stages, related to Figure 4.**

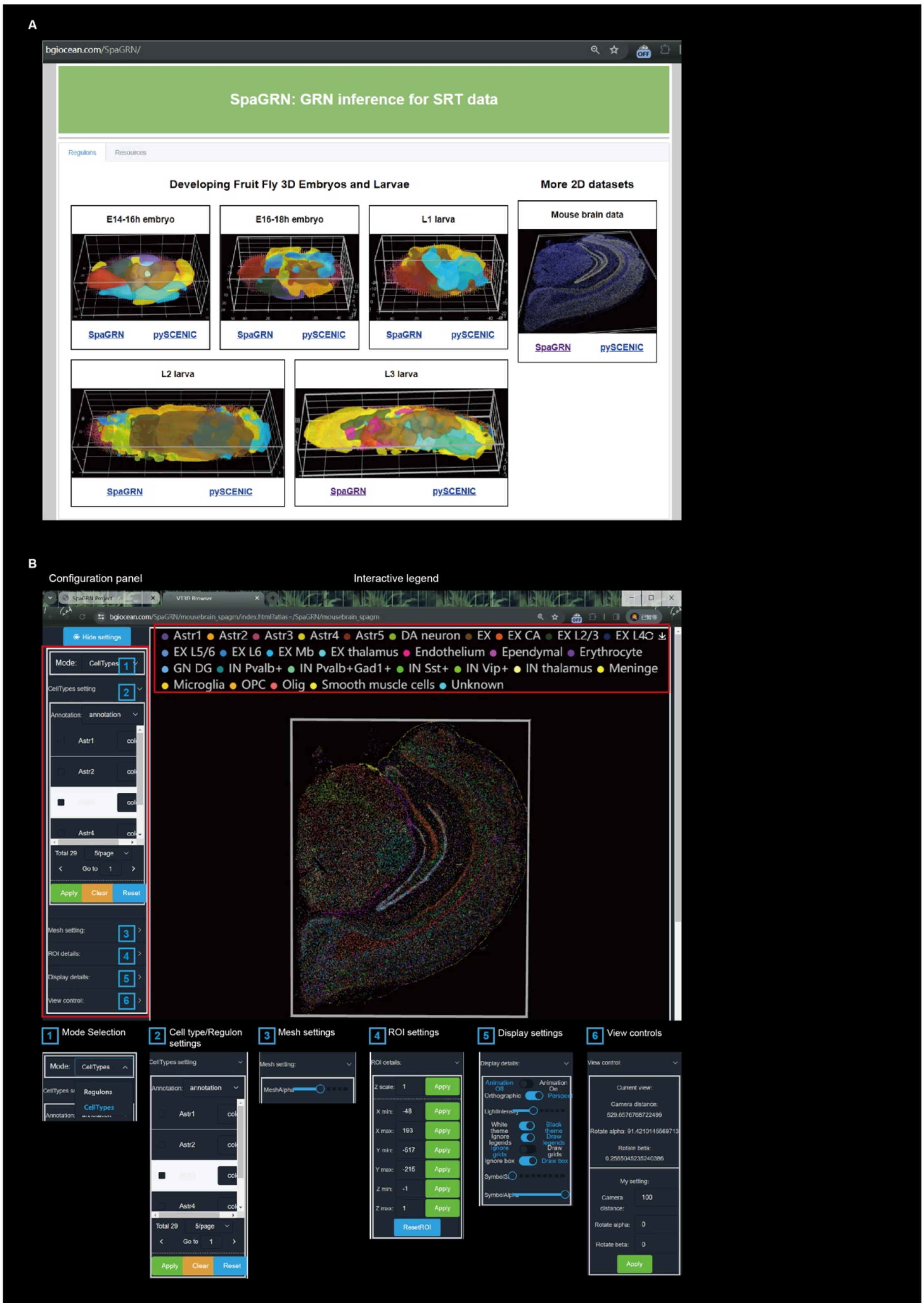


**Figure S6. 3D interactive visualization of SpaGRN results, related to Figures 3 and 4.**

**Table S1. Detail of datasets used in this study.**

| dataset | | sequencing platform | # slice | # cell or bin | # gene |
|---|---|---|---|---|---|
| adult mouse coronal hemibrain | | Stereo-seq | 1 | 50,140 | 25,879 |
| *Drosophila* embryos and larvae | E14-E16 | Stereo-seq | 13 | 15,295 | 13,668 |
| | E16-E18 | Stereo-seq | 11 | 14,634 | 12,850 |
| | L1 | Stereo-seq | 20 | 17,787 | 13,083 |
| | L2 | Stereo-seq | 19 | 64,658 | 14,270 |
| | L3 | Stereo-seq | 13 | 43,310 | 16,326 |

**Table S2. Statistics of computational performance on 3D fruit fly data, related to Figure 4.**

| dataset | tools | clock time (hour) | peak memory (Gb) |
|---|---|---|---|
| E14-16h | SpaGRN | 0.5 | 16.9 |
| | pySCENIC | 0.4 | 17.9 |
| E16-18h | SpaGRN | 0.3 | 16 |
| | pySCENIC | 0.3 | 16 |
| L1 | SpaGRN | 0.5 | 18 |
| | pySCENIC | 0.4 | 18.9 |
| L2 | SpaGRN | 4.8 | 55.6 |
| | pySCENIC | 1.4 | 56.8 |
| L3 | SpaGRN | 10.1 | 48.3 |
| | pySCENIC | 7.4 | 44.8 |

The GRN inferences were performed on a computer cluster with 20 Intel Core Processor (Broadwell, IBRS) CPU threads with 256 Gb of memory installed, and used the Linux operating system.
